# Virulence evolution of a parasite infecting male and female hosts

**DOI:** 10.1101/203687

**Authors:** Alison M. Wardlaw, Aneil F. Agrawal

**Affiliations:** University of Minnesota; University of Toronto

**Keywords:** evolution of virulence, non-random contact patterns, sex-specific disease

## Abstract

Parasites experience different tradeoffs between transmission and virulence in male and female hosts if the sexes vary in life history or disease-related traits. We determine the evolutionarily stable levels of exploitation by pathogens under two scenarios: an unconstrained pathogen that expresses different exploitation rates within each host type as well as a pathogen constrained to express the same exploitation rate in each sex. We show that an unconstrained horizontally-transmitted parasite evolves to express the same sex-specific exploitation rate within each sex as it would in a host population composed entirely of hosts with that sex’s resistance and intrinsic death rate. In contrast, the ESS exploitation rate of a constrained pathogen is affected by sex-differences in susceptibility and non-random contact patterns between host types that differ in resistance. As the amount of within-sex transmission increases, the ESS shifts closer to the optimum trait value in the more susceptible sex. Allowing for some degree of vertical transmission, the exploitation rate expressed in females (but not males) changes with contact pattern even in unconstrained pathogens. Differences in contact pattern and susceptibility play an important role in determining the ESS exploitation rate by shifting the reproductive value of each host type.

## Introduction

A pathogen’s fitness is determined by its ability to transmit to new hosts. Parasites with higher proliferation rates inside a host release more propagules for transmission but consume more host resources at the expense of increased disease-induced mortality (virulence; Anderson and May, 1982; Ewald, 1983; Alizon et al., 2009). The tradeoff between host mortality and transmission success affects parasite fitness: too low of an exploitation rate limits the number of new infections, but too high and the parasite kills its host before it has the opportunity for transmission. Selection therefore favours intermediate exploitation rates in classic one-host one-parasite models (May and Anderson, 1990; Frank, 1996).

Pathogens infecting more than one type of host face additional tradeoffs. The ability of a parasite to proliferate inside a host is affected by host immunity and available host resources, which can vary with condition, age, or sex (Day et al., 1991; Gaillard and Spinedi, 1998; Rice et al., 2000; Shankar, 2000; Bhaskaram, 2002; Field et al., 2002; Felix et al., 2012; Cousineau and Alizon, 2014, Table 1). Because realized virulence and transmission rates are the result of an interaction between host and parasite traits, a multihost parasite experiences a different tradeoff between host mortality and transmission success in each host type. The exploitation rate that maximizes parasite fitness in one host type may not be optimal in other host types. A pathogen in a heterogeneous host population can therefore adopt one of three strategies: 1) a single exploitation strategy, 2) alternative exploitation strategies maintained as a polymorphism, or 3) facultative expression of different exploitation strategies (Pfennig, 2001).

There are several general multihost parasite models that describe the evolution of a single exploitation rate (Gandon, 2004; Regoes et al., 2000; Osnas and Dobson, 2011; Williams, 2012; Cousineau and Alizon, 2014). These multihost models investigate the evolution of virulence using dynamical equations to describe the change in the number of susceptible and infected individuals of each host type over time. Gandon (2004) and Williams (2012) both analyzed multihost models where transmission and virulence were considered increasing functions of parasite exploitation rate and exploitation was scaled to capture different transmission and disease-induced mortality rates in each host. The evolutionarily stable level of exploitation depended on the relative abundance of host types and the ability of each type to transmit the infection. Gandon (2004) numerically investigated differential transmission between and within host types. At high within-type transmission rates, Gandon (2004) found evolutionary branching in parasite virulence, leading to the coexistence of different exploitation rates in the population. Williams (2012) developed a novel analytical way of thinking about multihost parasite evolution that also accounted for differences in susceptibility to infection (*i.e.* the likelihood that a given host type contracts the disease given contact with an infected individual). His model makes it easy to understand how between host interactions affect the importance of each host type to parasite fitness. The evolutionarily stable exploitation rate is a compromise between the ideal rates within each host type such that some host types are overexploited while others are underexploited.

If host types are affected differently by the same exploitation rate, differential disease-induced mortality across hosts could indicate that a parasite is constrained to express one trait. For example, Krist *et al.* (2003) found higher mortality rates in resource-limited snails infected with the trematode parasite *Microphallus* because the parasite did not adjust its development rate to optimally exploit low condition hosts. Similarly, a single exploitation strategy can have different effects in males and females because of differences in life history traits and immunocompetence (Cousineau and Alizon, 2014). Differential investment in reproduction in invertebrates can cause parasitized females to experience higher disease-induced mortality than males. For example, female damselflies infected by water mites suffered reduced mass at emergence and consequently decreased survivorship compared to males (Braune and Rolff, 2001). We can not be certain that the observed differences in virulence between host types in both of these examples is entirely due to inflexible parasite strategies. Because males and females and low and high condition hosts can also differ in immunocompetence, the host type with the higher survivorship could actually be suppressing an elevated type-specific parasite growth rate as opposed to experiencing the same exploitation rate as an overexploited host.

There are few documented examples of parasites facultatively expressing different strategies in different host types. Jokela et al. (1999) infected snails of high and low condition with two types of parasites independently, *Microphallus* and *Notocotylus gippyensis*. The authors observed higher mortality rates in parasitized snails from the no-food treatment, compared to the high food treatment, when infected with *N. gippyensis*. Food treatment did not affect disease-induced mortality when snails were infected with *Microphallus* indicating that the parasite adjusted its exploitation rate to host condition in order to ensure transmission. In another example, the *Ascogregarina culicis* parasite of mosquitoes (*Aedes aegypti*) has sex-specific strategies for releasing its infectious stages, called oocysts, to maximize overall transmission (Fellous and Koella, 2009). Even though there are more *A. culicis* oocysts in female mosquitos than in males at the pupal stage, the parasite releases a higher proportion of oocysts during male emergence. This maximizes transmission from male mosquitoes, while the parasite can wait to release oocysts during female ovipoistion when the mosquito will spread infectious stages to her offspring. From these examples it is apparent that evolving different strategies in heterogeneous host types could be evolutionarily advantageous. Untangling how the presence of other host types affects the strategies expressed requires further investigation.

Facultative expression of different exploitation strategies in each host type could evolve such that the parasite optimally exploits each host type or such that the parasite still under- or overexploits some host types. The optimal exploitation strategy in a host type is governed strictly by the within host tradeoffs a parasite faces between transmission and virulence. However, the presence of other host types could feedback and affect selection on transmission such that between host interactions favour type-specific exploitation strategies that are not the same as they would be in a homogeneous host population composed entirely of one host type. For example, it may be beneficial to overexploit some host types if it increases the probability of getting into a highly transmissive host type. The circumstances under which between host dynamics might feedback and affect facultative parasite expression are unknown. Modelling the expression of type-specific exploitation strategies could help us understand when epidemiological feedbacks are important and make predictions to be tested experimentally.

Between-host disease dynamics could be particularly important in determining the evolutionarily stable strategy when there is asymmetrical pathogen transmission. Gandon (2004) and Cousineau and Alizon (2014) both accounted for nonrandom contact patterns by symmetrically varying the amount of within and between host type transmission. An example of symmetrical contact patterns could occur in a monogamous mating system where each individual interacts primarily with a member of the opposite sex more than with members of the same sex. Alternatively, asymmetrical contact patterns could arise when males that compete for females have high within type transmission (*e.g.* if they have direct contact with one another while fighting for access to females), while female-to-female transmission rates are comparatively lower. In this scenario, males can be thought of as more valuable hosts to a pathogen than females. Varying the transmission route altogether, females may be more valuable hosts if there is vertical transmission in addition to horizontal transmission of the disease. Because asymmetrical transmission could change a pathogen's relative fitness out of each host type, we might expect different evolutionary outcomes for a constrained parasite than observed in previous multihost models.

Our model builds on that of Williams (2012), incorporating heterogeneous virulence responses in the form of resistance, and adding nonrandom transmission between and among host types. This framework allows us to tease apart the role of resistance, susceptibility, and transmission patterns in determining the evolutionarily stable parasite strategy and understand how these disease-related traits affect each other’s relative importance. We also explicitly model facultative expression of exploitation rates by assuming two independent pathogen traits, exploitation rate in each host type.

Overall, we find that when there are differences in resistance between host types, differential susceptibility to disease can change evolutionary outcomes for a parasite constrained to express a single exploitation strategy, especially at high within-host type transmission rates. In comparison, an unconstrained parasite evolves to express the same exploitation rate in each host type as it would in a homogeneous population of that host type, regardless of differences in susceptibility or contact pattern. Treating the two host types as males and females and allowing for some vertical transmission, an unconstrained parasite will evolve an evolutionarily stable exploitation strategy in females that changes with contact pattern to take advantage of the alternative transmission route from mother to offspring. Our results show that transmission patterns and differential susceptibility can have important effects on virulence evolution in heterogeneous host populations.

## Model Setup

We use a system of differential equations to describe a susceptible-infected compartmental model with two host types, which we refer to as males and females throughout. Apart from the sections incorporating reproductive input and vertical transmission, the model can be applied to other kinds of two host-type systems where individuals do not transition between types. In the general model we assume new uninfected individuals enter the population at a constant immigration rate, *θ*, a fraction *σ* of which are males and the remaining (1 − *σ*) are females. Uninfected individuals of sex *i* become infected with the disease at the rate *η*_*i*_*S*_*i*_(*p*_*ii*_*β*_*i*_*I*_*i*_ + *p*_*ji*_*β*_*j*_*I*_*j*_) where *η*_*i*_ is the sex-specific probability of contracting the disease given contact with an infected individual, and *p*_*ii*_*β*_*i*_*I*_*i*_ + *p*_*ji*_*β*_*j*_*I*_*j*_ is the force of infection on host type *i*. The force of infection depends on the number of infected males and females (*I*_*f*_ and *I*_*m*_) and the parasite’s transmission rate out of each (*β*), modified by the probability of encounter (or transmission opportunity) between infected type *j* and uninfected type *i* individuals (*p*_*ji*_). Finally, uninfected individuals of type *i* die at the natural host-mortality rate, *μ*_*i*_, and infected individuals suffer from additional disease-induced mortality (virulence, *α*_*i*_). Transmission rates and virulence both depend on the parasite's intrinsic exploitation rate, *ϵ*_*i*_, inside its host. A host type with some level of resistance against the disease, *ρ*_*i*_, can reduce the parasite’s effective exploitation rate, consequently reducing disease-induced mortality and transmission rates (see equation [2]). The disease dynamics are captured by the following set of differential equations.

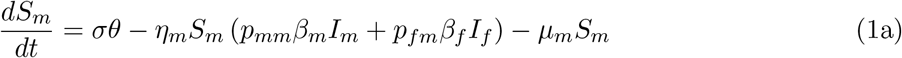

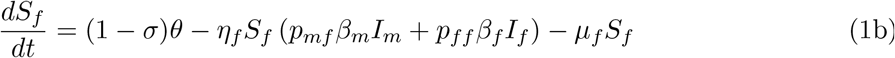

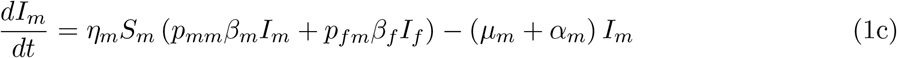

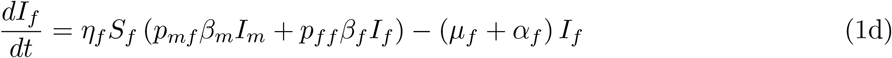

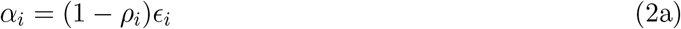

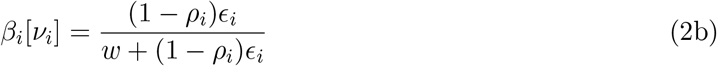

For a parasite with a given exploitation strategy, the population will reach a steady state where the number of individuals leaving each uninfected and infected class is equal to the number of individuals entering that class. We can write the steady state number of uninfected and infected males as:

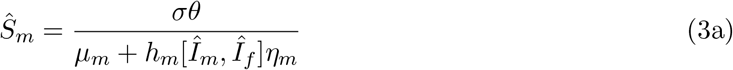

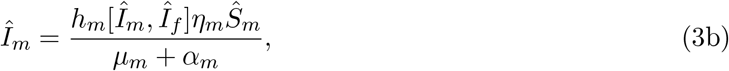

where *h*_*i*_[*Î*_*i*_, *Î*_*j*_] = *p*_*ii*_*β*_*i*_*Î*_*i*_ + *p*_*ji*_*β*_*j*_*Î*_*j*_ is the equilibrium force of infection on host type *i*. *Ŝ*_*f*_ and *Î*_*f*_ are defined analogously but *σ* is replaced with 1 − *σ*.

We can model the evolution of the parasite using an adaptive dynamics approach, in which a mutant parasite with a slightly different exploitation rate is introduced into a resident host-parasite system at equilibrium to determine if it has higher fitness and will replace the resident. The successive invasion of mutants can lead to an evolutionary endpoint which is an evolutionary stable strategy if it is uninvadable by nearby mutants (Otto and Day, 2007). To determine if a mutant will invade, we augment the model to include a mutant allele and perform a linear stability analysis. Using tildes to denote variables and parameters pertaining to the mutant, the dynamics of the mutant are given by equation (4).

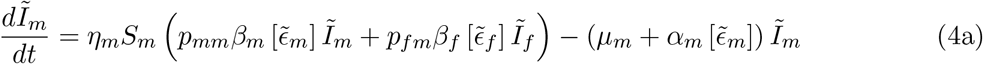

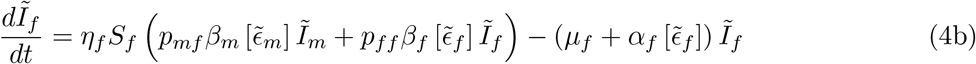

**Table 1:**
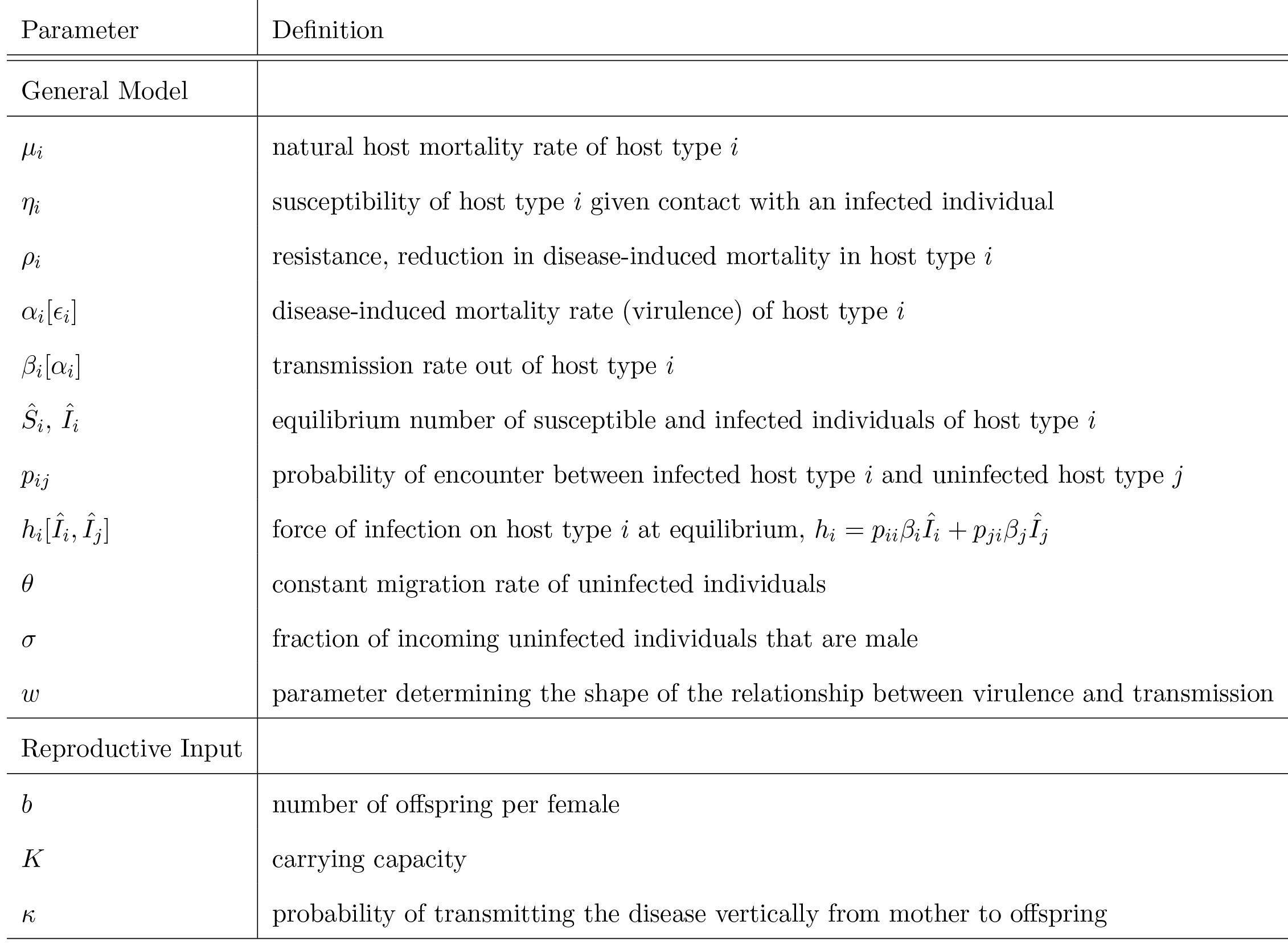
List of parameters and their definition

| Parameter | Definition |
| --- | --- |
| General Model |  |
| $\mu_i$ | natural host mortality rate of host type $i$ |
| $\eta_i$ | susceptibility of host type $i$ given contact with an infected individual |
| $\rho_i$ | resistance, reduction in disease-induced mortality in host type $i$ |
| $\alpha_i[\epsilon_i]$ | disease-induced mortality rate (virulence) of host type $i$ |
| $\beta_i[\alpha_i]$ | transmission rate out of host type $i$ |
| $\hat{S}_i, \hat{I}_i$ | equilibrium number of susceptible and infected individuals of host type $i$ |
| $p_{ij}$ | probability of encounter between infected host type $i$ and uninfected host type $j$ |
| $h_i[\hat{I}_i, \hat{I}_j]$ | force of infection on host type $i$ at equilibrium, $h_i = p_{ii}\beta_i\hat{I}_i + p_{ji}\beta_j\hat{I}_j$ |
| $\theta$ | constant migration rate of uninfected individuals |
| $\sigma$ | fraction of incoming uninfected individuals that are male |
| $w$ | parameter determining the shape of the relationship between virulence and transmission |
| Reproductive Input |  |
| $b$ | number of offspring per female |
| $K$ | carrying capacity |
| $\kappa$ | probability of transmitting the disease vertically from mother to offspring |

We are interested in the stability of the equilibrium where the mutant allele is absent, *i.e*. 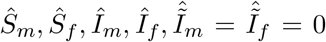. Assuming the resident allele is stable before the introduction of the mutant, the mutant will invade if the leading eigenvalue, 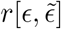, of the mutant transition matrix **J**_*Mut*_ is greater than zero.

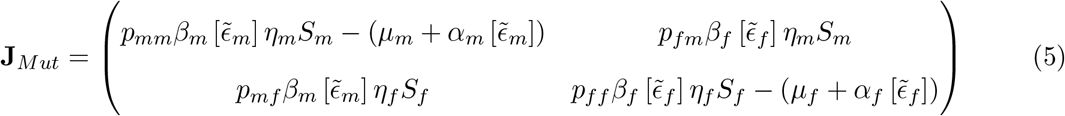

### Homogeneous Host Population

For a parasite with a single exploitation strategy (*ϵ*_*m*_ = *ϵ*_*f*_ = *ϵ*), the direction of evolution (*i.e*. successive invasions) of the parasite trait is given by the fitness gradient, defined as 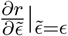. In a homogeneous host population (*i.e*. males and females are phenotypically equivalent) the fitness gradient equals 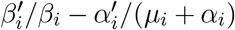. Because this term arises repeatedly, it is convenient to define 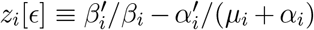 following Williams (2012). The first term in *z*_*i*_[*ϵ*] represents the relative transmission benefit of an increase in exploitation rate, while the second term represents the relative virulence cost. If the fitness gradient is positive (negative), the parasite is under(over) exploiting its host and a mutant parasite with a higher (lower) exploitation rate will invade. When the fitness gradient equals zero, the pathogen is at an evolutionary endpoint, which is evolutionarily stable if it is a fitness maximum and any nearby mutants move toward that level of exploitation (convergence stability). The evolutionarily stable strategy (ESS) of a constrained parasite in a homogeneous host population composed entirely of host type *i* is 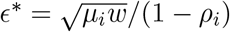 (Otto and Day, 2007).

### Heterogeneous Host Population

A parasite in a heterogeneous host population could be (1) constrained to express the same exploitation rate in both sexes, or (2) free to facultatively express different exploitation rates in males and females, *ϵ*_*m*_ and *ϵ*_*f*_, respectively, where *ϵ*_*m*_ and *ϵ*_*f*_ are modelled as two independently evolving traits.

#### Two Parasite Traits

We determined the evolutionary endpoints for a facultative parasite by performing a multivariate invasion analysis (Otto and Day, 2007). In the unconstrained case, the evolutionarily stable strategy must simultaneously satisfy equations (6a) and (6b),

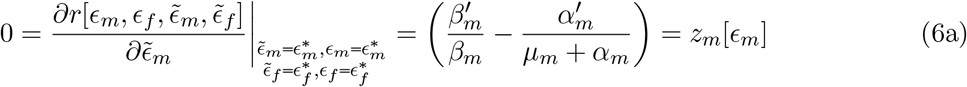

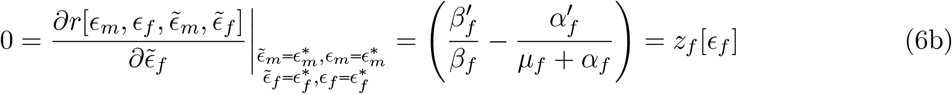

whose solution is 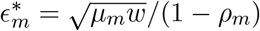 and 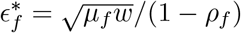.

We see that a parasite in a heterogenous host population that can infect both sexes is selected to express the same exploitation rate in a given sex as it would if it were in a population composed entirely of hosts with that sex’s phenotype with respect to intrinsic death rate and resistance. The relative abundance of each sex and differential disease transmission between sexes does not affect the strategy a pathogen should adopt in its host. In other words, epidemiological feedbacks do not play a role in the evolutionary outcome when parasite fitness only depends on optimizing transmission out of its host (Mideo et al., 2008).

#### One Parasite Trait

Constraining *ϵ*_*m*_ = *ϵ*_*f*_ = *ϵ*, we examine this ESS exploitation rate in comparison to the unconstrained case where 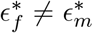. When transmission rate depends only on the parasite’s exploitation rate inside its host *i.e*. transmission between and among the sexes is random, Williams (2012) found that the evolutionarily stable exploitation rate satisfies equation (7) where the *z*_*i*_[*ϵ*] are weighted by each host type’s relative contribution to the force of infection

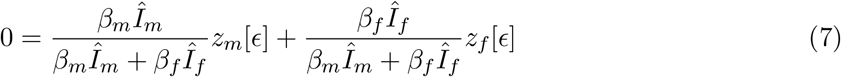

These weightings however, assume that an infected host interacts with any type of uninfected host in the same way with regards to disease spread. We relaxed that assumption, allowing for differential transmission between and among sexes. Our result, given by equation (8), has a similar form but the *z*_*i*_[*ϵ*] are weighted by that sex’s relative contribution to the force of infection on the opposite sex.

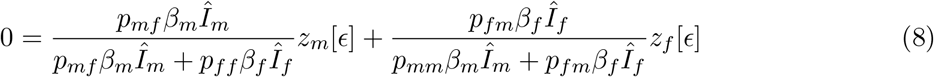

The *p*_*ij*_ terms represent the probability of the disease spreading from an infected host of type *i* to an uninfected host of type *j*. Even though the *z*_*i*_[*ϵ*] weightings do not include the relative contribution of within sex transmission to the force of infection, within sex transmission still affects the ESS determination. For example, decreasing female to female transmission increases the relative contribution of between sex transmission to the force of infection on females. *z*_*m*_[*ϵ*] is weighted more heavily than when there is random transmission so the ESS will shift closer to 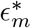. It is easier to interpret equation (8) if we rewrite it in terms of the number of infected individuals of each sex and their relative contribution to the parasite’s long-term growth rate. Respectively, these factors are captured by the right and left eigenvectors, 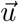 and 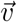, of **J**_*Mut*_ (Otto and Day, 2007). Writing the fitness gradient as shown in equation (9), we see how the stable class distribution 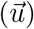 and class reproductive values 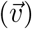 affect the evolution of the parasite trait.

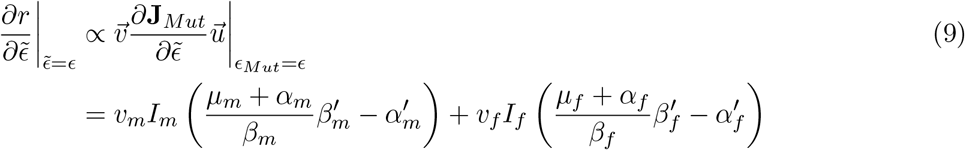

The transmission benefit accompanying a small increase in exploitation rate depends on the rate infection ends (*μ*_*i*_ + *α*_*i*_) in a host relative to the transmission rate (*β*_*i*_) out of that host. In other words, the faster the infection ends, the more beneficial it is to increase transmission to new hosts but there is less to be gained when transmission out of host type *i* is already high. Increasing disease-induced mortality carries a direct cost. The tradeoff between transmission and virulence within each sex is affected by between host disease dynamics. The number of infected males and females and their reproductive value to the parasite (*υ*_*m*_ and *υ*_*f*_) determines how the pathogen balances the tradeoff it faces in each sex. The reproductive values not only depend on the intrinsic transmission rate out of each sex, *β*_*i*_, but also on transmission opportunity to uninfected hosts (eq. 10). For example, the contribution of an infected male to the population growth rate of the parasite depends on its transmission rate to uninfected males and females, *p*_*mm*_*β*_*m*_*η*_*m*_*S*_*m*_ and *p*_*mf*_*β*_*m*_*η*_*f*_*S*_*f*_, respectively, and the future reproductive value of the newly infected individuals of each sex, *υ*_*m*_ and *υ*_*f*_. The duration of infection 1/(*μ* + *α*_*i*_) also affects sex *i*’s reproductive value.

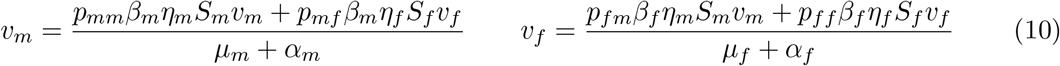

## Results

To get a better understanding of equation (9), we plot the constrained ESS for different contact patterns when the source of new uninfected individuals is constant immigration (fig.1). We focus on how host traits directly related to disease, *e.g.* resistance and susceptibility, affect the evolutionarily stable parasite strategy relative to values of 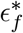 and 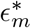. We therefore assume that the natural host mortality rate is the same in males and females (*μ*_*m*_ = *μ*_*f*_ = *μ*).

### Single differences between host types

#### Susceptibility

If susceptibility is the only difference between the sexes, then the population behaves as a homogeneous host population (Williams, 2012). In this case, *α*_*m*_[*ϵ*] = *α*_*f*_ [*ϵ*] = *α* and *β*_*m*_[*ϵ*] = *β*_*f*_ [*ϵ*] = *β*. ((*μ* + *α*)/*β*)*β*′ − *α*′ can be factored out of equation (9) and solved for the one host one parasite solution 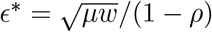 (Otto and Day, 2007).

#### Resistance

To isolate the effects of differences in resistance between the sexes, we assume equal susceptibility (*η*_*m*_ = *η*_*f*_ = *η*) and equal numbers of each sex entering the population (*σ* = 0.5). When transmission is symmetrical, the analytical solution for the constrained ESS, *ϵ*\*, is given by 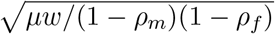, the geometric mean of 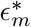 and 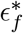. This solution does not depend on contact pattern. Regardless of the amount of within sex transmission the density of uninfected males and females is equal (not shown). Effectively, the parasite strategy reflects the fact that it is equally likely to be transmitted into either sex. Assuming males are less resistant than females, we plotted the unchanging ESS against increasing within sex transmission rates (black solid line, fig.1) in order to facilitate comparison with the ESS when there are multiple differences between the sexes.

### Multiple differences between host types

When the sexes differ in both susceptibility and resistance, contact pattern does affect the evolutionarily stable exploitation rate. Compared to when only resistance is different between sexes, differential susceptibility (*η*_*m*_ ≠ *η*_*f*_) shifts the ESS closer to that of the more susceptible sex (black dashed and dotted dashed lines, fig.1). The relative importance of susceptibility depends on the amount of within sex transmission (fig. 1). When there is only between type transmission, *i.e. p*_*mf*_ = *p*_*fm*_ = 1, the ESS equals that of *η*_*m*_ = *η*_*f*_. The parasite has to pass through each sex sequentially and is therefore equally likely to be transmitted to uninfected males or uninfected females. As the amount of within sex transmission increases, the ESS moves closer to 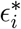 of the more susceptible sex. Once the parasite gets into that sex, it is much more likely to be transmitted within it so being closer to 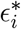 has a long term evolutionary advantage. More precisely, the parasite has a higher reproductive value in the more susceptible sex (fig. 2), making the transmission virulence tradeoff in that sex more important in the ESS determination.

At high within sex transmission rates we observe evolutionary branching because the host population is effectively two subpopulations with no disease transmission between them. When susceptibility is equal evolutionary branching is observed at within sex transmission rates greater than 0.94. Differences in susceptibility between the sexes increase the within sex transmission rate above which evolutionary branching occurs (*p*_*ii*_ > 0.99) because the parasite is specialized on the more susceptible sex. Since most of its transmission opportunities are in the more susceptible sex, a single specialized strategy is still a fitness maximum unless between sex transmission is so rare that males and females represent distinct subpopulations.

### Asymmetrical disease transmission

In the previous section we assumed contact patterns were symmetrical, meaning that an increase in male to male transmission rates was accompanied by an increase in female to female transmission rates. However, in their efforts to obtain mates, males may interact with each other and with females more than females interact with one another. This would result in asymmetrical opportunity for disease transmission. Asymmetrical transmission patterns can lead to changes in the ESS with contact pattern even in the absence of differences in susceptibility between the sexes (fig. 3a). Figure 3a shows the constrained and unconstrained ESS as a function of increasing transmission into males. We first consider the case where transmission probability out of either sex into males is the same and total transmission probability out of each sex is constant, so that increasing transmission probability to males is accompanied by reduced transmission to females from both sexes (*p*_*mm*_ = *p*_*fm*_ = *δ*; *p*_*mf*_ = *p*_*ff*_ = 1 − *δ*) As expected, the ESS is closer to 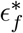 when transmission is mostly into females and closer to 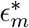 when transmission is mostly into males (dashed green line, fig. 3a). Reproductive values in males and females are similar (fig. 3b) but there are many infected individuals of the sex disproportionately contracting the disease, skewing the ESS towards that 288 sex’s unconstrained optimum.

**Figure 1:**
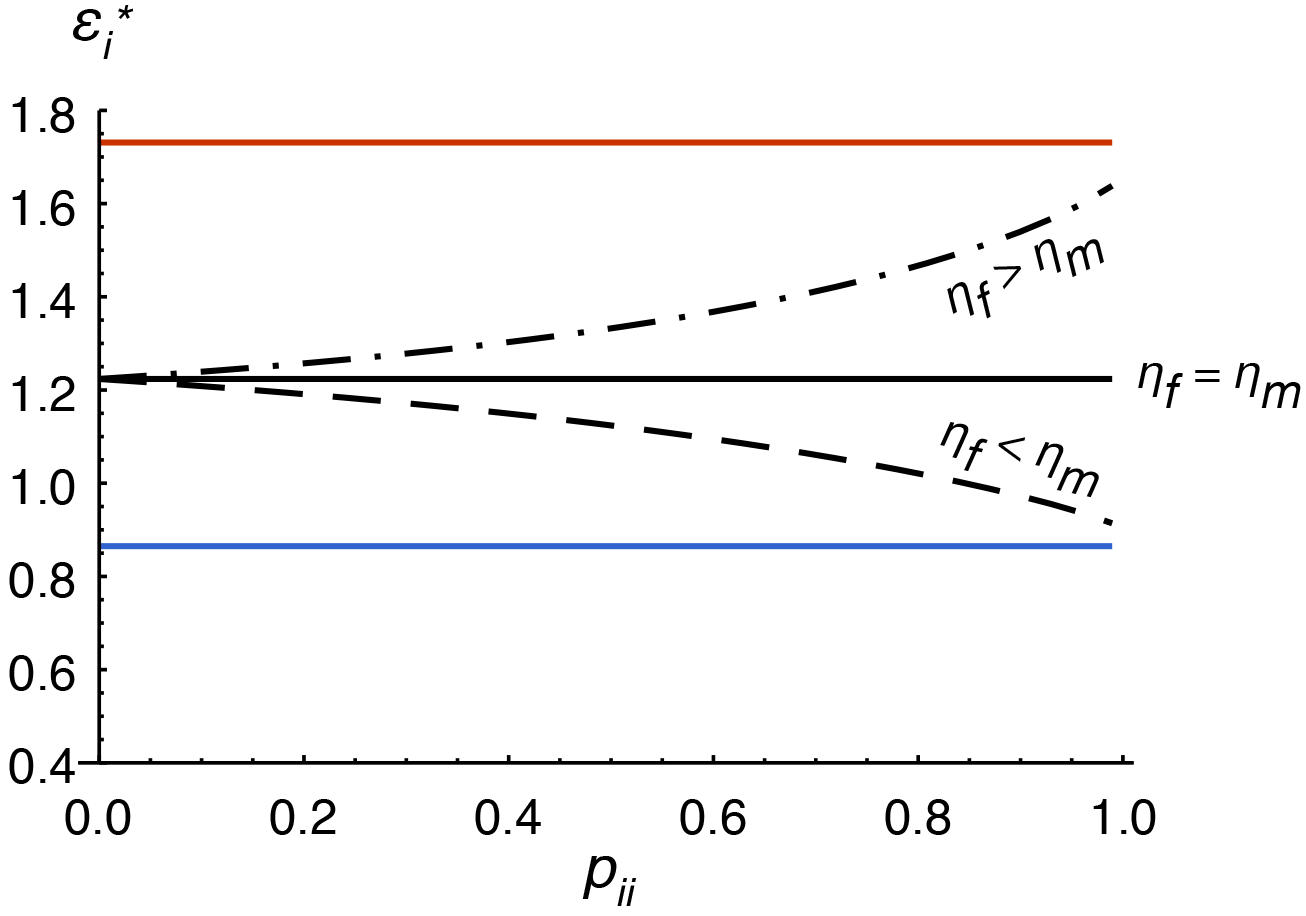
The evolutionarily stable strategy (ESS) of a parasite expressing the same exploitation rate in both sexes changes with contact pattern when there are differences in susceptibility between the sexes. Transmission is symmetrical (*p*_*ij*_ = *p*_*ji*_ and *p*_*ii*_ = *p*_*jj*_) and transmission coefficients out of each sex sum to one (*p*_*ij*_+*p*_*ii*_ = 1) such that as within-sex transmission increases moving from left to right across the *x*-axis, between-sex transmission decreases. A parasite that facultatively expresses different exploitation strategies in males and females has ESS exploitation rates 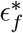 (red) and 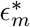 (blue), which do not change with differential susceptibility or contact pattern. The constrained ESS also does not change with contact pattern when susceptibility is equal (*η*_*f*_ = *η*_*m*_, black solid line) but shifts closer to 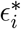 of the more susceptible sex as within sex transmission increases (*η*_*f*_ > *η*_*m*_, black dashed dotted line; *η*_*f*_ < *η*_*m*_, black dashed line). The source of new uninfected individuals is constant immigration, *i.e. θ* = 75. Though the figure goes to *p*_*ii*_ = 1 for illustrative purposes, evolutionary branching occurs below this level (*e.g. p*_*ii*_ > 0.94, with the exact value depending on the parameters)

**Figure 2:**
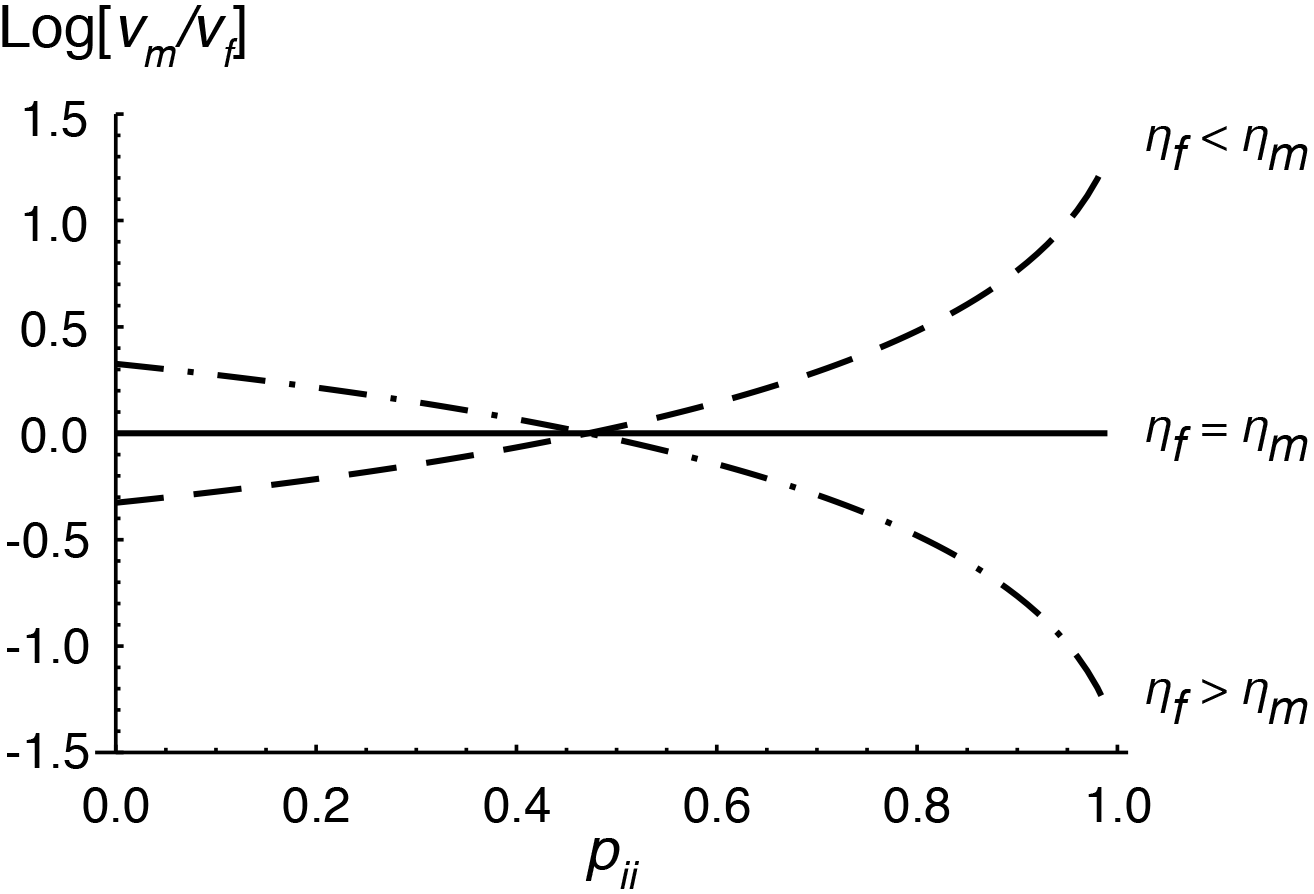
A constrained parasite has higher reproductive values in the more susceptible sex as within-sex transmission increases. The class reproductive values (*υ*_*m*_ and *υ*_*f*_) represent the contribution of each sex to future population growth of the parasite. Males and females contribute equally to parasite population growth when susceptibility is equal (*η*_*f*_ = *η*_*m*_, black solid line). When there is differential susceptibility (*η*_*f*_ > *η*_*m*_, black dashed dotted line; *η*_*f*_ < *η*_*m*_, black dashed line), the parasite has a higher reproductive value in the less susceptible sex if there is more between-sex transmission or in the more susceptible sex if there is more within-sex transmission. The source of new unifected individuals is constant immigration, *i.e. θ* = 75 and the *x*-axis is the same as in figure 1.

When we hold transmission out of females constant at *p*_*ff*_ = *p*_*fm*_ = 0.5 and consider changes in transmission probability out of males (*p*_*mm*_ = *δ*; *p*_*mf*_ = 1 − *δ*) we might still expect the ESS to be closer to 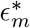 at high male to male contact rates. However, the ESS is closer to 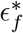 even when transmission into males is high (solid green line, fig. 3a). The reproductive value in females is higher than that in males (fig.3b) because females can transmit the disease to either sex. Furthermore, regardless of the extent to which males transmit the disease amongst themselves (and not to females), there is always some disease transmission among females, maintaining enough infected females to be important in the ESS determination. This is not true for low input rates (*θ* < 55) where there can be so few infected females that the ESS is closer to 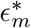 even though the pathogen’s reproductive value is lower in males at high *δ* (not shown).

**Figure 3:**
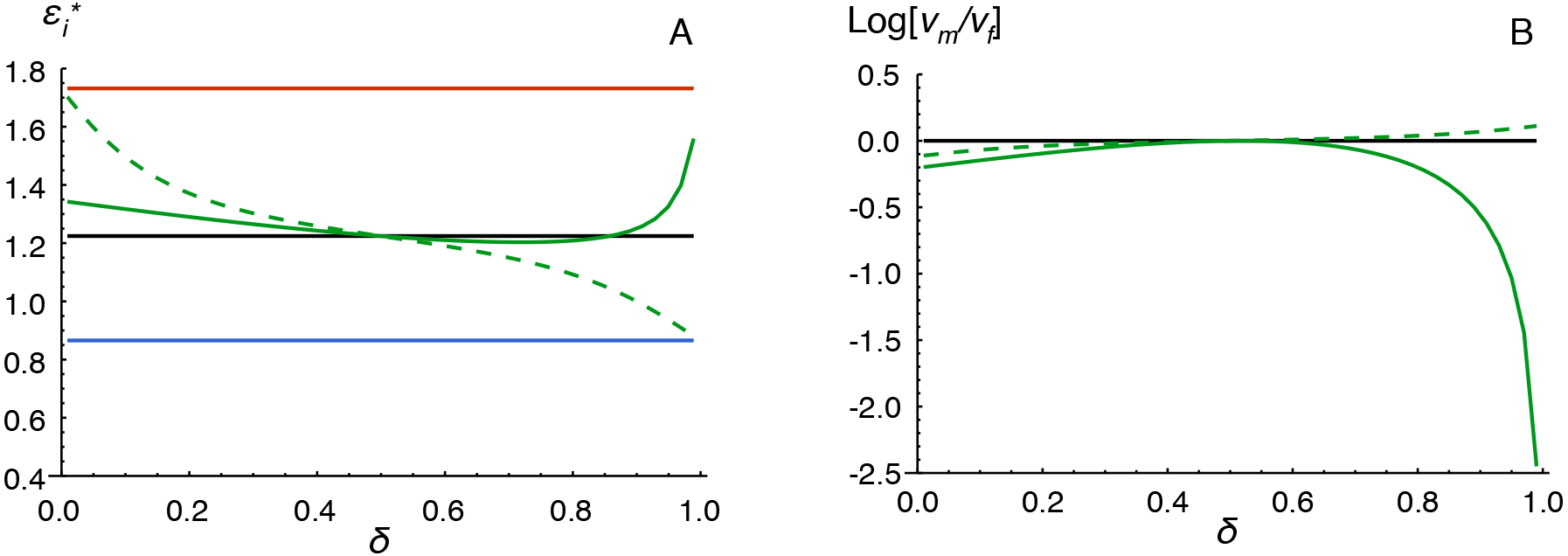
(A) Asymmetrical transmission patterns lead to changes in the ESS in the absence of differential susceptibility, *i.e. η*_*f*_ = *η*_*m*_. Two scenarios are depicted: transmission into males from females and other males increases moving from left to right across the *x*-axis (*p*_*mm*_ = *p*_*fm*_ = *δ*; *p*_*mf*_ = *p*_*ff*_ = 1 − *δ*, dashed green line) or male to male transmission increases while transmission out of females remains random (*p*_*mm*_ = *δ*; *p*_*mf*_ = 1 − *δ*; *p*_*ff*_ = *p*_*fm*_ = 0.5, solid green line). The two trait ESS values, 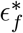 and 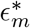, are shown as the red and blue solid lines, respectively. (B) Relative class reproductive values for a one trait parasite. The source of new uninfected individuals is constant immigration, *i.e. θ* = 75.

### Source of new uninfected individuals

If new uninfected individuals in the population arise through density dependent births, ecological feedbacks could affect the evolution of the parasite. We consider two additional types of input: female dominant density dependent reproduction (eq.11a) and density dependent reproduction where both males and females are important for the production of offspring (eq.11b). These input terms, which are a function of the number of uninfected and infected males and females, replace the constant migration rate *θ* in equation (1). Here, *b* represents the number of offspring per female, following (Lindström and Kokko, 1998) and (*S*_*f*_ + *I*_*f*_)(*S*_*m*_ + *I*_*m*_)/*N* is the harmonic birth function (Caswell, 1989). Density dependence is incorporated through logistic growth where *K* is the carrying capacity of the population.

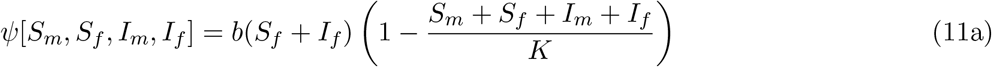

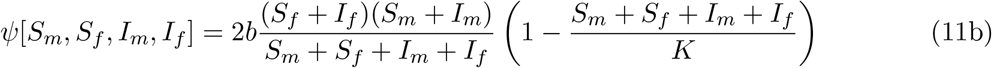

The source of new uninfected individuals into the population has an important effect on the evolutionarily stable exploitation rate when males and females are not equally susceptible to disease (*η*_*m*_ ≠ *η*_*f*_). For density dependent births where both sexes are important to reproduction, the constrained ESS is almost equal to that of a parasite in a population of similar size experiencing constant immigration (fig. 4b). When population growth is based on female dominant density dependent births, the ESS is closer to the unconstrained female optimum than when both sexes are equally important or when there is constant input of uninfected individuals (fig.4a). This pattern arises because of changes in the density dependent birth rate and overall population size with contact pattern (not shown). Smaller population sizes with differential susceptibility to disease show greater divergence away from the equal susceptibility ESS than larger ones at high within sex transmission rates (not shown). Under female dominant reproduction when females are more susceptible, birth rate and population size decrease with increasing within sex transmission rates and the ESS diverges further away from the *η*_*f*_ = *η*_*m*_ ESS (towards 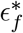, dashed dotted black line, fig. 4a). When males are more susceptible, females suffer less from the disease at high within sex transmission rates. Population size is larger and the ESS shifts towards the *η*_*f*_ = *η*_*m*_ ESS (dashed black line, fig. 4a). Regardless of the input type, ecological feedbacks in the density dependent models can affect the ESS of a constrained parasite. However, the ESS of the unconstrained parasite remains at the optimum in males and females.

**Figure 4:**
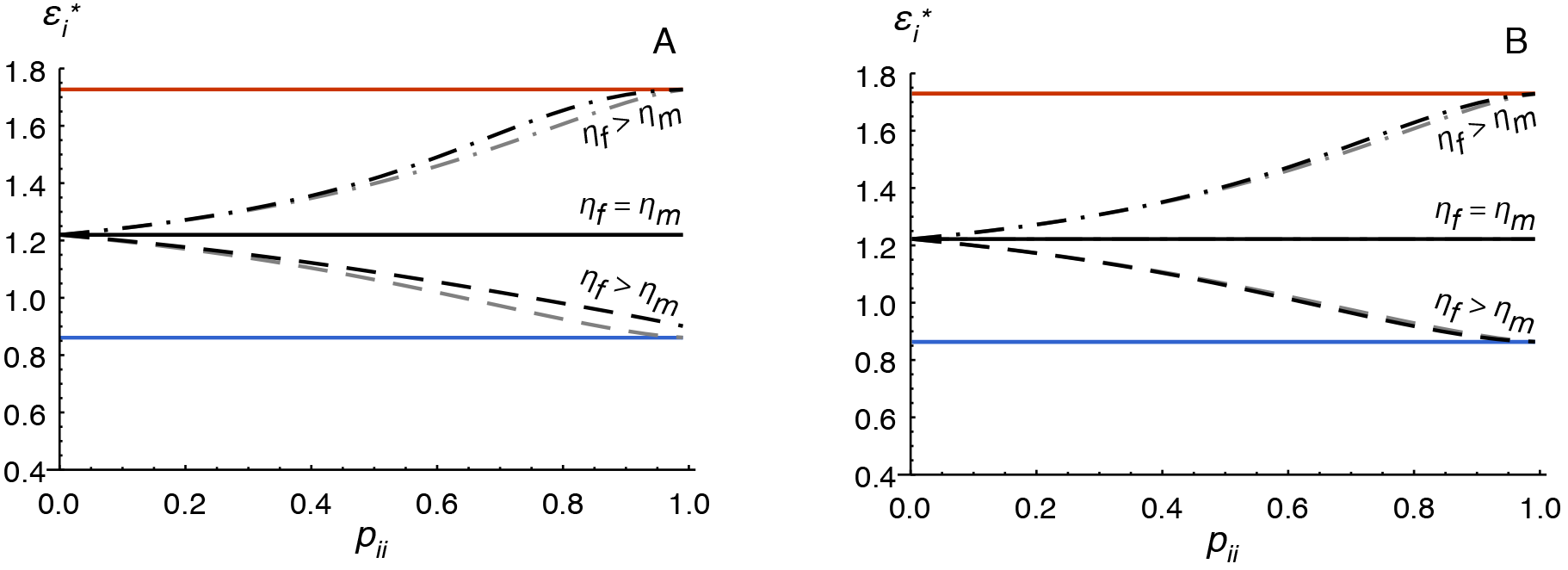
The source of new uninfected individuals can affect how differential susceptibility skews the ESS towards 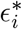 of the more susceptible sex. Density dependent births where females are more important to reproduction (A, black lines) shift the ESS closer to 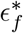 compared to when both sexes are important to reproduction (B, black lines) or when there is constant immigration (both panels, grey lines). A parasite that facultatively expresses different exploitation strategies in males and females has ESS exploitation rates 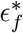 (red) and 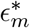 (blue), which do not change with differential susceptibility or contact pattern. *η*_*f*_ = *η*_*m*_, solid lines; *η*_*f*_ > *η*_*m*_, dashed dotted lines; *η*_*f*_ < *η*_*m*_, dashed lines. The *x*-axis is the same as in figure 1.

### Horizontal versus Vertical Transmission

We incorporate vertical transmission into the female dominant density dependent model by allowing infected females to give birth to uninfected offspring with probability 1 − *κ* and infected offspring with probability *κ*, where *κ* is *not* a function of the parasite’s exploitation rate. Allowing some vertical transmission has several effects. The ESS exploitation rate of females in the unconstrained model is now sensitive to the extent of within- versus between-sex transmission. The ESS in females changes with contact pattern if the disease is vertically transmitted because its long term transmission success depends on whether it is being transmitted to males or females. When the parasite is only horizontally transmitted, all that matters is maximizing transmission out of its current host (see eq. 6). With vertical transmission (*κ* = 0.5, fig.5a), females offer two modes of transmission and are more valuable hosts. If there is only between-sex transmission, a parasite is better off increasing the duration of infection in its female host than transmitting to males (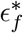 is lower than 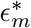). As within-sex transmission increases, female transmission is increasingly into other females. Higher exploitation rates in females ensure the parasite is transmitted to as many females as possible (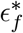 is higher than 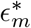). As with a horizontally transmitted parasite, the ESS of the constrained model is bounded by the male and female ESS exploitation rates of the unconstrained model.

Vertical transmission also causes the ESS exploitation rate in females of an unconstrained pathogen to be sensitive to differences in susceptibility between the sexes. This is easiest to see at low rates of vertical transmission (*κ* = 0.1) where 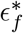 changes less drastically with increasing within-sex transmission rates (fig. 5b).

**Figure 5:**
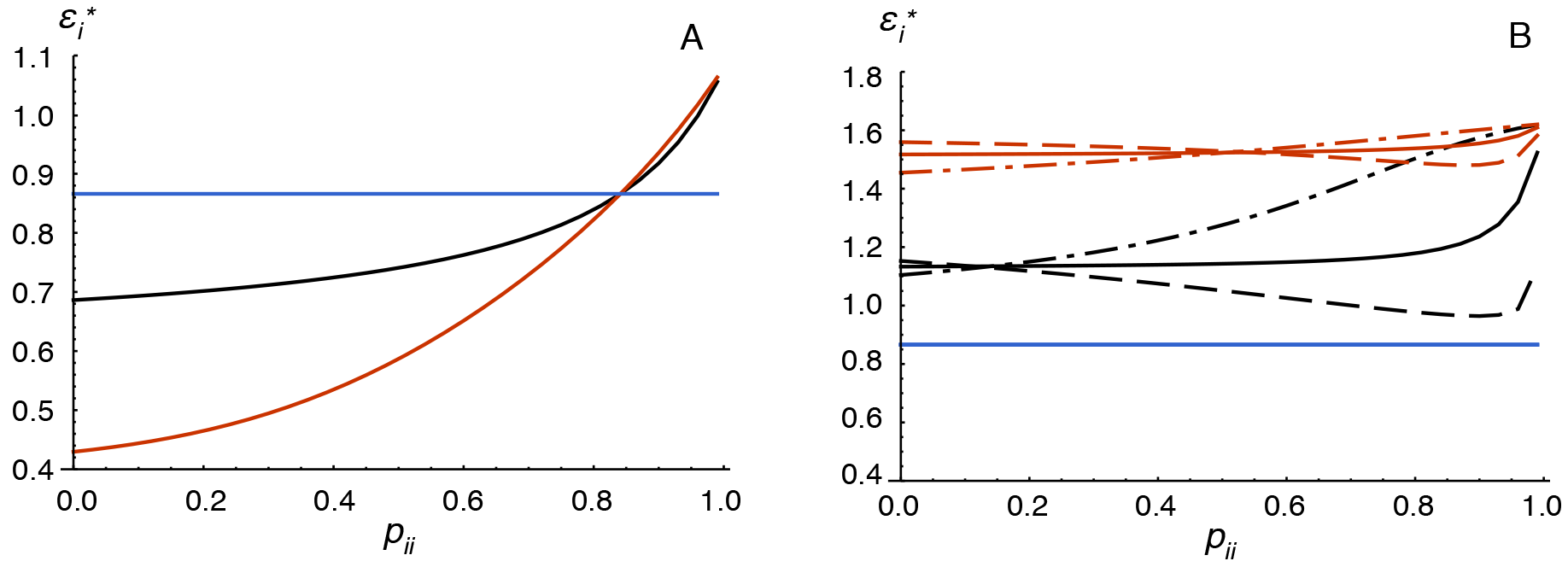
For an unconstrained parasite, the exploitation rate expressed in females changes with contact pattern when there is vertical transmission. (A) The probability of vertical transmission is *κ* = 0.5. 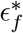 increases with increasing within-sex transmission (solid red line) while 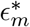 does not (solid blue line). The constrained parasite strategy (solid black line) is always bounded by 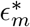 and 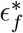. Only the equal susceptibility case is shown for clarity. (B) The probability of vertical transmission is *κ* = 0.1. 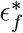 changes with contact pattern to a lesser extent. Solid lines, *η*_*f*_ = *η*_*m*_; dashed dotted lines *η*_*f*_ > *η*_*m*_; dashed lines *η*_*f*_ < *η*_*m*_. The x-axis is the same as in figure 1.

## Discussion

We expanded the multihost parasite model developed by (Williams, 2012) to incorporate differential transmission between two host types. For horizontally transmitted parasites, transmission patterns did not change the evolutionary outcome when only one of resistance or susceptibility were different between the sexes. When both were different, the relative importance of susceptibility in determining the evolutionary stable exploitation strategy is greater when the probability of within-sex transmission is large relative to between-sex transmission. We also explicitly modelled an unconstrained parasite, that is, a parasite that can express different exploitation rates in each sex. We found that an unconstrained parasite evolves to express the same exploitation rate in a given sex as it would in a homogeneous population of that sex, regardless of susceptibility or contact pattern, unless there is vertical transmission of the disease. Thus, differential transmission patterns create changes in disease dynamics that feed back to the affect the constrained ESS, but do not feed back to affect the ESS in the unconstrained model unless females are intrinsically more valuable hosts than males because they are capable of vertical transmission.

Even though an unconstrained parasite expresses different evolutionarily stable exploitation rates in males and females, transmission out of and virulence in each sex is the same because 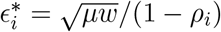 and *α*_*i*_ = (1 = *ρ*_*i*_)*ϵ*_i_. This observation arises because we allow exploitation rate in males and females to be independent traits. It has been established that between-host disease dynamics do not affect a parasite’s strategy for maximizing transmission out of its host unless parasite life history traits depend on uninfected and infected host densities (Mideo et al., 2008). Alternatively, 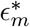 and 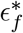 will only differ from the homogeneous host population expectation if there is some correlation between traits (Gandon, 2004). In our model, between-host disease dynamics do not feed back - regardless of how relative host densities change with differential susceptibility and transmission patterns - to affect parasite proliferation within a host when there is only horizontal transmission.

Epidemiological feedbacks do, however, affect the ESS of a horizontally transmitted one-trait parasite. Because the parasite is constrained to express one exploitation rate in both males and females, virulence in males is not equal to virulence in females. The difference in virulence, and consequently transmission, means that changes in susceptibility or transmission patterns differentially affect the parasite’s ability to transmit from each sex. Though the changes in between-host disease dynamics are not feeding back to affect the tradeoffs within a host, they still affect the relative contribution of transmission from each sex to overall transmission success. The tradeoff between virulence and transmission faced in each sex is therefore weighted differently depending on susceptibility to disease and the amount of between- and among-sex transmission.

When only resistance is different between the sexes, 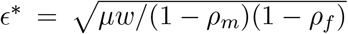. We can compare our results to those of Cousineau and Alizon (2014) who studied the evolution of virulence in sexually dimorphic hosts. Cousineau and Alizon (2014) investigated varying levels of sexual dimorphism in tolerance and resistance, separately, for only between-sex transmission, random transmission, and mostly within-sex transmission. Host tolerance decreased disease-induced mortality rates while host resistance decreased disease-induced mortality and transmission rates. For maximal levels of sexual dimorphism in their model, one sex had no resistance to disease and the other sex’s resistance varied from zero to one. They found the rate of increase in the ESS with increasing resistance varied depending on the transmission pattern. In our model, the transmission pattern does not affect the ESS when susceptibility in both sexes is equal. Their model differs from ours in several ways, including how resistance is incorporated into the model. More importantly, Cousineau and Alizon (2014) assume constant population size and an equal sex ratio. If we also assume constant and equal numbers of each sex, the ESS does change with contact pattern under equal susceptibility (*η*_*m*_ = *η*_*f*_), moving closer to the male optimum with increasing within-sex transmission. Because males are overexploited, there are higher transmission rates out of males and hence more male to male transmission. In our model increased transmission among males is balanced by increased male mortality rates resulting in the same ESS as with random transmission. Holding population size and sex ratio constant, high disease-induced mortality in males means dying infected males are replaced with uninfected males. The parasite has more transmission opportunities to uninfected males and the ESS shifts closer to 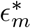.

Contact patterns determine if a constrained parasite will be a specialist or a generalist given sex-specific susceptibility. High within-sex transmission causes the parasite to specialize on the more susceptible sex while high between-sex transmission drives generalism. Regoes et al. (2000) investigated a parasite with a free-living stage that traded off virulence in one host type against that in the other (*e.g. ϵ*_*i*_*ϵ*_*j*_ = *c*). They found that specialism arose when the shape of the tradeoff curve was convex enough that the cost of switching host types was high. While we did not explicitly model a virulence tradeoff, the pathogen effectively faces a tradeoff between host types because of sexual dimorphism in resistance. When specialized on one sex because of high transmission opportunity within that sex, the pathogen will over or under exploit its new host type if it is transmitted between sexes. The cost of switching increases with within-sex transmission rates until it is so high that evolutionary branching occurs and two constrained exploitation rates are maintained in the population as a polymorphism.

Vertical transmission selects for less virulent pathogens because the parasite cannot be transmitted from mother to offspring if the mother dies of disease-related causes before reproducing (Yamamura, 1993). Our results are consistent with theory on vertical transmission in host pathogen systems (Lipsitch et al., 1995). The constrained exploitation rate and the exploitation rate expressed in females by an unconstrained parasite are both lower than predicted by a female dominant density dependent model with only horizontal transmission (fig. 5). The extent to which the parasite evolves lower exploitation rates depends on the amount of vertical transmission. Since the disease cannot be transmitted from father to offspring in our model, the male optimum is the same as predicted by equation (6a) and does not change with contact pattern. At high vertical transmission rates, an unchanging 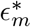 can result in a switch in which sex is over exploited and which is underexploited (fig. 5a). This switch could affect the epidemiological dynamics and have an important impact on predicting and managing the disease in male and female patients.

Considering resistance, susceptibility, and transmission patterns in isolation can over or underestimate evolutionary stable levels of virulence. Careful examination of host disease traits and contact patterns will lead to the best understanding of disease dynamics and evolutionary endpoints. This is particularly important because different host types will differ not only in immune responses such as resistance, but also in susceptibility to infection and the ways that different host types interact with one another to cause disease transmission.

